# Identifying dynamic reproducible brain states using a predictive modelling approach

**DOI:** 10.1101/2022.10.14.512147

**Authors:** D O’Connor, C Horien, F Mandino, RT Constable

## Abstract

Conceptually brain states reflect some combination of the internal mental process of a person, and the influence of their external environment. Importantly, for neuroimaging, brain states may impact brain-behavior modeling of a person’s traits, which should be independent of moment-to-moment changes in behavior. A common way to measure both brain states and traits is to use functional connectivity based on functional MRI data. Brain states can fluctuate in time periods shorter than a typical fMRI scan, and a family of methods called dynamic functional connectivity analyses, have been developed to capture these short time estimates of brain states. There has been a rise in the use of dynamic functional connectivity in order to find temporally specific spatial patterns of connectivity which reflect brain states, that can yield further insight into traits and behaviors. It has previously been shown that brain state can be manipulated through the use of continuous performance tasks that put the brain in a particular configuration while the task is performed. Here we focus on moment-to-moment changes in brain state and test the hypothesis that there are particular brain-states that maximize brain-trait modeling performance. We use a regression-based brain-behavior modelling framework, Connectome-based Predictive Modelling, allied to a resample aggregating approach, to identify behavior and trait related short time brain states, as represented by dynamic functional connectivity maps. We find that there is not a particular brain state that is optimal for trait-based prediction, and drawing data from across the scan is better. We also find that this not the case for in-magnet behavioral prediction where more isolated and temporally specific parts of the scan session are better for building predictive models of behavior. The resample aggregated dynamic functional connectivity models of behavior replicated within sample using unseen HCP data. The modelling framework also showed success in the estimating variance behavior in the ABCD dataset when using data from that dataset. The method detailed here may prove useful for both the study of behaviorally related brain states, and for short time predictive modelling.

## Introduction

Measuring brain function using fMRI and functional connectivity (FC) has yielded increased insight into whole brain functional topology and the relationship between brain function and behavior [1], [2]. FC has been shown to be individually unique [3] and has been used to investigate and derive reproducible brain-behavior relationships [4], [5]. The majority of FC based studies aggregate information across an entire four-dimensional scan into a one- or two-dimensional “static” measure of FC, for example a graph theory metric [6], or an adjacency matrix [7]. While these metrics have provided insight into brain-behavior relationships, they are, by design, insensitive to events which vary on shorter time scales.. This can obscure potentially cognitively relevant patterns present in the moment-to-moment data. Indeed, it has been shown that FC topology varies on timescales much shorter than a typical scan length [8], [9], and the characteristics of such variations may reflect inter-subject trait differences [10], [11]. Studies incorporating temporal variation in FC into their analyses are on the rise [12]. Such investigations use what has been described as time varying or dynamic functional connectivity (dFC).

It has been shown that the task a participant is engaged in during data acquisition can be determined from time windows as short as tens of seconds [13]. Variations at this time-scale have been tied to changes in vigilance and cognitive engagement [14];they can be used to predict human error [15], to assess cognitive decline in an elderly population [10], and to investigate dFC reorganization in healthy and diseased brains, e.g. between typical and schizophrenic participants [16]. Methods for estimating dFC have emerged in order to encapsulate as much data from one individual’s scan as possible [12], [17]–[19]. With these dFC methods the goal is often to find brain “states”, which are temporal snapshots of connectivity thought to be relevant to a behavior, or a trait. One of the more common approaches involves using sliding window estimates of FC, which allow calculation of correlations between timeseries at shorter timeframes or windows [20]. Other methods for deriving short time FC include phase synchrony connectivity [10], [21], tapered sliding windows, spatial distances, and temporal ICA, a partial list is detailed here [22]. Phase synchrony connectivity is of particular interest, as it allows derivation of “instantaneous” connectivity, which can be estimated at the temporal resolution of the data acquisition.

Once calculated, these windowed estimates of FC from different time points can be clustered into groups. The prototypical approach by Allen et al. used *k*-means based clustering to accomplish this [23]. The clusters or groups of dFC reflect different functional connectivity configurations, often referred to as brain states. Following this, dwell time within states and state-switching frequency can be calculated. These metrics have been considered as indicators of trait based differences [19].Other brain state detection methodologies include Hidden Markov Modelling [11], [24], change point estimates [25], [26], and dictionary learning based “windowless” connectivity [27]. There are also methods for defining temporally overlapping brain states. The common hypothesis in these approaches is that there is a reproducible set of common spatiotemporal patterns that the brain transitions through during a given scan, which can provide insight into the cognitive processes ongoing during data acquisition.

While dFC allows the identification of temporally discrete FC, these metrics are based on fewer data points, and thus have lower SNR, poorer reliability, and are more susceptible to typical noise processes, overall [18]. Whilst changes in resting dFC may be the result of unconstrained cognition fluctuations [28], sampling error, motion and arousal can also affect these changes [29]. Validation of dFC patterns is difficult, as we often lack a ground truth for what spurs changes in dFC. One approach to statistical validation of dFC changes is null modelling. Many dFC studies attempt to construct null spaces, simulating random data from real covariance matrices and power spectra [29], [30]. Using these methods, one can potentially distinguish cognitively relevant dFC from noise patterns. Further, since the majority of dFC is performed on resting state fMRI data [12], another way to mitigate a lack of ground truth is to use task-based fMRI (for a detailed review on the task/rest dichotomy, in the context of dFC, see Gonzalez-Castillo and Bandettini [31]). Studies assessing noise dominance on dFC focus mostly on resting state fMRI [29]; however, it has been argued that integrating task and rest dFC can improve confidence in the relationships estimated [19]. More reproducible, and synchronous, changes in FC can be driven by a common task. Both null modelling and task based dFC have the potential to validate dFC approaches.

Model-based approaches have are frequently used to capture the relationship between static FC and behavior/traits. These methods allow for statistical validation via model building and testing. Even with some statistical validation, such as null modelling, it is important to test the generalization of any statistical relationship derived. One modelling method which has shown great promise in FC-based modelling is connectome-based predictive modelling (CPM) [4]. CPM has been used to successfully predict attention [5], abstinence from drug use [32], obesity [33], and ADHD/autism related traits [34]. CPM studies typically involve the application of models to unseen data, to ensure a level of reproducibility in the estimated relationship. A similarly rigorous approach can be used to assess the reproducibility of dFC based models. Predictive modelling-based approaches have been used with dFC, though mostly classification-based, for example task classification [13], and disease classification (schizophrenia and bipolar disorder) [35]. Regression-based approaches to dFC brain-behavior modelling are few. Fong et al. used dFC estimates to model attention performance [36], though still aggregating data across whole scans into the model. Nonetheless, the success of both these studies suggests that regression-based approaches, such as CPM, have the potential to both identify brain states from short time dFC and link them to specific behaviors or traits.

In this study, we use a combination of a regression model-based approach, CPM, and phase synchrony based dFC to identify specific temporal periods, or brain states, that are related to a working memory-based trait and short time-varying behavior. We contrast the predictive power of rest and task-based fMRI, as well as trait and behavior, and show that task-based prediction of behavior may be more optimal for identifying temporally unique brain states. We assess the models derived using cross validation, within sample testing and out of sample testing, showing good generalizability of these brain state models within sample. Our study offers a framework for identifying and validating behavior related brain states using predictive modelling.

## Methods

### Datasets

Data from the Human Connectome Project (HCP), specifically the S900 dataset, [37] and the Adolescent Brain Cognitive Development (ABCD) [38], from the Fast-track Imaging Data Release (year one arm), were used. From these datasets, N=827 participants from the HCP, ages 21-35, N=1668 ABCD participants from the ABCD, ages 9-11, were included. Inclusion criteria for both data sets were based on the availability of preprocessed T_1_-weighted images, N-Back task fMRI scans, ePrime files containing task trial information, resting state fMRI scans and, in the case of HCP, fluid intelligence (fIQ) measures. See Rapuano et al. for more about the inclusion criteria for ABCD [39].

For the HCP dataset, fIQ was measured using a 24-item version of the Penn Progressive Matrices assessment, scores ranged from 4-24, with a mean ± standard deviation (SD) of 16.78 ± 4.7 [40]. The score corresponds to the number of correct responses.

HCP MRI data were acquired on a 3T Siemens Skyra. The fMRI scans were collected using a slice-accelerated, multiband, gradient-echo, echo planar imaging (EPI) sequence (TR = 720ms, TE = 33.1ms, flip angle = 52°, resolution = 2.0mm^3^, multiband factor =8). The T_1_-weighted structural scans were collected using a Magnetization Prepared Rapid Gradient Echo (MPRAGE) sequence (TR = 2400ms, TE = 2.14ms, TI = 1000ms, resolution = 0.7mm^3^) [41].

ABCD MRI data used in this study were acquired using Siemens Prisma or Philips scanners with a 32-channel head coil. Detailed acquisition parameters have been previously described in the literature [38]. Scan sessions included a high-resolution T_1_-weighted scan, resting-state fMRI, and task-based fMRI. Functional images for both scanners were collected using an accelerated multiband echo-planar imaging sequence (TR = 800 ms, TE = 30 ms, flip angle = 52°, resolution = 2.4 mm^3^, multiband slice acceleration factor = 6). The Siemens T_1_-weighted structural scans were collected using Magnetization Prepared Rapid Gradient Echo (MPRAGE) sequence (TR = 2500ms, TE = 2.88ms, TI = 1060ms, resolution = 1mm^3^). The Philips T_1_-weighted structural scans were collected using Magnetization Prepared Rapid Gradient Echo (MPRAGE) sequence (TR = 6.3ms, TE = 2.9ms, TI = 1060ms, resolution = 1mm^3^).

### Preprocessing

For the HCP, the HCP minimal preprocessing pipeline was used on these data [42], which includes artifact removal, motion correction, and registration to MNI space. All subsequent preprocessing was performed in BioImage Suite [43] and included standard preprocessing procedures [3], i) with removal of motion-related components of the signal, ii) regression of mean time courses in white matter, cerebrospinal fluid, and gray matter, iii) removal of the linear trend, iv) low-pass filtering.

ABCD data were preprocessed using BioImage Suite [43] with an approach described in detail elsewhere [39], [44], [45]. T_1_-weighted anatomical images were skull stripped using optiBET [46], and non-linearly registered to MNI stereotaxic space using B-spline free form deformation. Functional images were realigned to correct for motion, and registered to MNI space. Further preprocessing was performed as above for the HCP data, including i) removal of motion-related components of the signal, ii) regression of mean time courses in white matter, cerebrospinal fluid, and gray matter, iii) removal of the linear trend, iv) low-pass filtering.

### Connectivity Measures

For both datasets, ROI time series were generated from all fMRI data using the Shen 268 atlas [47] (which defines 268 cortical and subcortical nodes). Dynamic connectivity matrices were generated from the ROI timeseries using phase coherence-based connectivity. This method has been demonstrated previously [10], [21], and has been found to yield similar results to sliding window correlation For both datasets, ROI time series were generated from all fMRI data using the Shen atlas [47] (which defines 268 cortical and subcortical nodes). Dynamic connectivity matrices were generated from the ROI timeseries using phase coherence-based connectivity. This method has demonstrated previously [10], [21], and has been found to yield similar results to sliding window correlation [48], though with the advantage of being able to estimate connectivity at single timepoint resolution. In brief, phase coherence dFC can be calculated by:

1. Applying the Hilbert Transform to the parcellated fMRI timeseries, resulting in two temporal estimates of the time series; magnitude and phase. The magnitude time series are discarded.
2. The phase time series was used to calculate instantaneous connectivity by taking the pairwise differences of phases for different ROIs at the timepoint level.
3. The cosine function was applied to these phase differences, yielding one connectivity matrix of n regions x n regions for each timepoint (268×268 for the Shen atlas ROIs).

The matrix values are bounded between 1 and −1; where 1 indicates maximally aligned phases, 0 indicates unaligned phases, and −1 anti-aligned phases. In this study we generated dFC estimates at the instantaneous level, and also generated connectivity estimates for varied window lengths, by taking the mean of the instantaneous connectivity across the window. The window lengths we decided depending on the target variable. For fIQ prediction, window lengths of 1, 5, 10, 30 60, 120, 240, 300, 350, and the full scan were used. For N Back response time prediction, window lengths of 1, 9, 15, 21, and 29 were used. Odd numbers of frames were chosen for response time so that windowed estimates of connectivity could be centered around the time of a given response time, drawing even numbers of frames from before and after the event. Twenty-nine timepoints was the largest window possible, whereby the start/end of the scan was not reached.

As counterpoints to the standard approach of creating windowed estimates of dFC, i.e., from fMRI volumes in the order they were acquired, we also generated dFC from within scan shuffled task data. That is, a random vector of the same length as the scan was applied for each scan of every participant, rearranging the order of the fMRI volumes. This resulted in data which was misaligned across participants, with respect to time and task structure. We also generated dFC from a given participants resting state data. Each of these different data types was used in the modelling process.

In all cases, the connectivity matrices generated are undirected, and thus symmetric around the diagonal. As such, 35,778 edges from each matrix were used as features for modelling.

### Modeling Protocol

For our modelling framework we chose to use CPM. CPM was performed as in [4], with the dFC matrices as the explanatory variable, and either a trait (fIQ) or behavior (N-Back response time, RT) as the target variable, with one exception; partial correlation was used at the feature selection step [49]. In full, the CPM process was as follows:

1. Across the training set, correlate each element in the FC matrix, often referred to as an edge, (predictive variables) with fIQ (target variable). In this work, partial correlation was used. First, mean frame-to-frame displacement in the fMRI scan is regressed out of both the edge values and fIQ. Then Pearson’s correlation is computed from the residuals.
2. Select positively correlated edges with, where p < 0.1.
3. For each participant or scan, sum the connectivity scores for all selected edges.
4. Fit a linear model, without regularization, between the sum of connectivity scores and fIQ summary score.
5. Apply the model to unseen participants in the testing set and estimate performance.

In this study we performed split half cross validation (CV) as an initial test of model performance for both fIQ and RT. For RT we applied further testing. We aggregated features and model parameters across all the CV folds and created a mean model and feature vector, as described previously in O’Connor et al [50]. As will be described in more detail below, this mean model and feature vector was tested in sample, and out of sample.

#### Dataset handling

Following participant exclusion as in the Datasets section above the HCP dataset contained 827 subjects, and the ABCD dataset contained 1,668 subjects. These datasets were split into subsets based on our analytic approach. The HCP data was split into two subsamples; a sample of 500, to be used for cross validation-based assessment of the modelling approach, and a separate evaluation test set of 327 for evaluating resample aggregated models generated from first sample. The separate evaluation test set was used exclusively to test within-sample performance, and none of the participants were included in model training.

There were 4 different task block orders for the 1668 subjects in the ABCD dataset. In order to facilitate cross subject comparison four separate subsamples were created with homogenous task block ordering. This resulted in four subsamples with 426, 393, 402, and 447 subjects respectively, labelled ABCD samples 1-4. The ABCD data was used both for CV based estimation of RT prediction, and for testing HCP models.

#### Model Training

In the HCP 500 sample, split half CV was performed using dFC to predict both fIQ and RT. Following CV, resample aggregated models were generated for timepoints which showed the highest levels of predictive performance. For each of the ABCD groups, split half CV was performed using dFC to predict RT. In each case 50 iterations of split half CV was performed. In the case of the resample aggregated models this yielded 100 resamples across which to aggregate feature occurrence and model parameters.

#### Model Testing

The HCP test sample is used to assess the within-sample performance of all resample aggregated models generated, and the ABCD data is used to assess out-of-sample performance of these same models. In within-sample testing each model is tested on 50 random subsamples of ~180 participants from the HCP test sample. In out of sample testing the models were evaluated on 50 random subsamples of 300 participants from each of ABCD samples.

In generating the resample aggregated models, the same approach as detailed in O’Connor et al. was used [50]. In short, a feature vector is created, in which each element corresponds to the frequency with which a given feature passes feature selection, across resampling. This allows the definition of a minimum frequency threshold, and the exclusion less frequently occurring features from being included in the model. The resample aggregated models are first tested including every feature which occurred at least once in any single resample. Following this, the feature vector is thresholded. First, to include features which occur in 10% or more resamples, then 20% or more, rising in 10% increments, to those that occur in 90% or more of resamples. For each threshold, the resample aggregated models’ performance is assessed within- and out-of-sample for each data split, as described above. In each case the models were assessed with different levels of thresholding applied to the feature occurrence vector. Thresholds were applied so the features were included when they occurred at least once across resamples, and then in 10% steps up to features that occurred in greater than or equal to 90% of resamples. This framework yields 50 measures of performance for each model, for each threshold, within-sample, and 50 measures of performance for each model, for each threshold, for each sample of ABCD data, out-of-sample. Model performance was assessed using to R^2^.

All analyses were performed in python using custom scripts. These scripts have been made publicly available, see section on data and code availability. Figures were generated in python using plotly, nilearn, matplotlib [51] and seaborn [52].

## Results

### Prediction of trait

We first attempted to predict a trait, fIQ, from dFC (Figure 1, top panel). Window lengths ranging from 1 timepoint (instantaneous connectivity) to the full scan length were used in prediction. As a point of comparison, both shuffled task data and resting state data were used to predict fIQ as well. The first comparison (shuffled task data) was made to ascertain whether the temporal ordering of the data matters. The second comparison (resting data) was made to ascertain whether data without task driven cognitive processes, but similar noise processes, have predictive power. As shown with the task data, it is possible to predict fIQ from relatively little data (i.e. explaining 2-3% of variance, with a window length of 30 timepoints, on par with expected effect sizes of static resting state FC [53]), though the more data included, the better the performance. Interestingly, in the range of 10-120 timepoint window lengths, shuffled working memory tended to outperform temporally ordered data, with its median performance explaining 1-3% more variance in the scores. This is notable given the highest median performance for task-based prediction of fIQ in this study was around 9% variance explained. While it is possible to predict fIQ from relatively small amounts of data, better performance is achieved when volumes are drawn randomly from across the whole scan, or when more data are included. This suggests that there are not optimal, temporally specific brain states in isolated parts of the scan for predicting fIQ. Both task-based prediction methods outperformed resting state for all window lengths.

**Figure 1.**
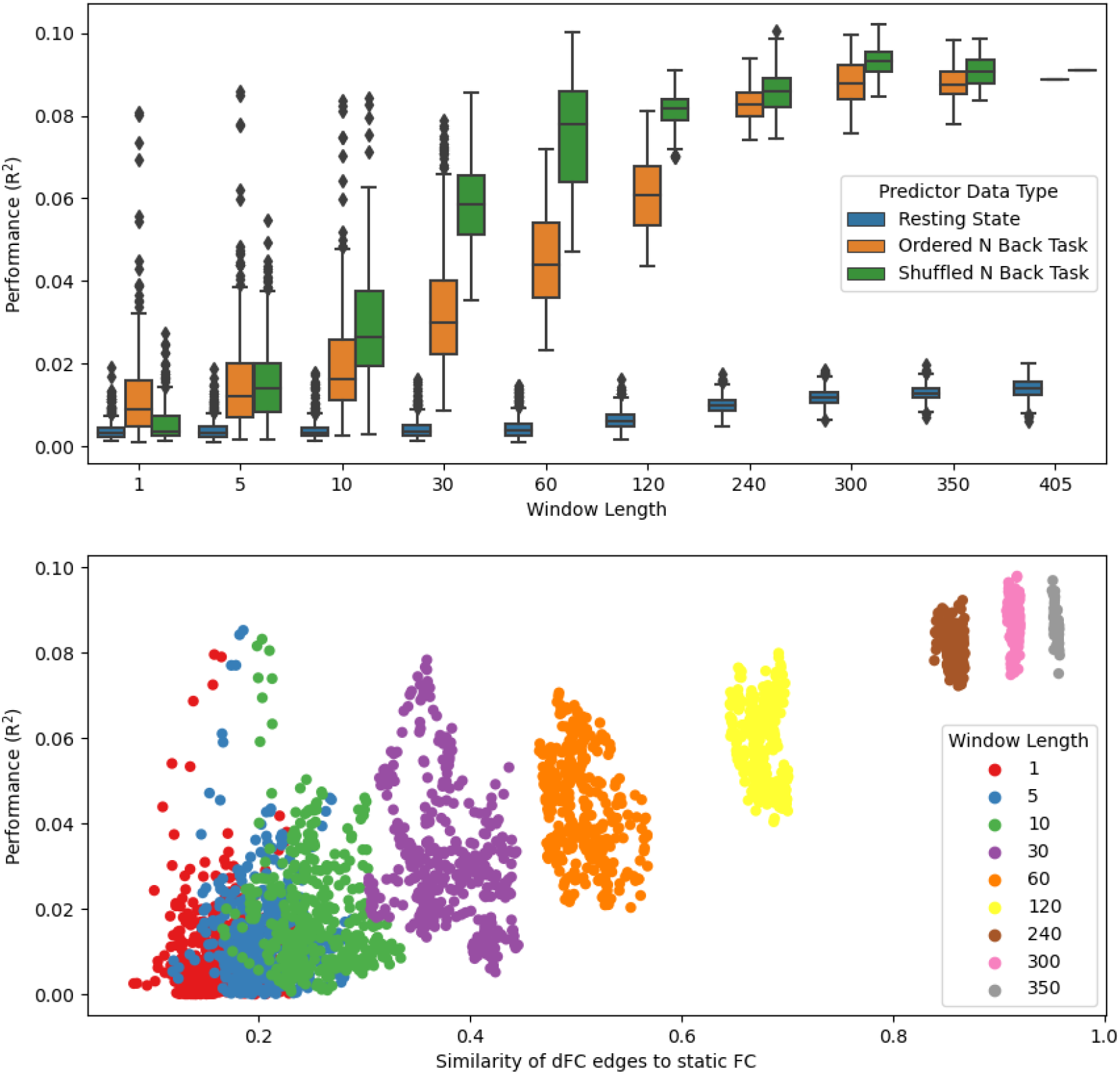
Performance of dFC models across time windows, for prediction of fIQ. Upper panel shows the predictive performance (y axis) of resting state dFC (blue), shuffled task dFC (green), and ordered task dFC (orange), for different window lengths (x axis). Lower panel shows the performance of ordered task dFC (y axis) versus the similarity of short time dFC features to features derived at the static level. Window lengths are labelled by color, see legend.

While the performance of short window dFC never rose to the levels of static FC, we were also curious if the features being selected in short time dFC models were distinct from those selected by the static model, which may suggest that different parts of the scan reflect different brain states. To assess this, we contrasted the features selected in each of the dFC models with those selected in the static model. This was done be generating mean feature vectors for each timepoint prediction, for each window length in the ordered task data, and correlating them with the mean feature vector from the static FC modelling approach (Figure 1, lower panel). When the features extracted at the dynamic level tended to be more similar to the static features (the window lengths are longer), the better they performed. This suggested that short time scale predictive features provide little benefit in modelling time-invariant traits.

### Prediction of Time-varying Behavior

#### Cross validation: dFC across the scan

Following the above analysis, and acknowledging that traits do not vary within scan, we next investigated trait measures that potentially do exhibit unique spatiotemporal FC patterns in a time-varying manner. We tested the hypothesis that real-time behavioral measures, which are related to traits, and which may vary across a scan, may be reflected in dFC models. As such, we attempted to build models of dFC and response time to the N Back task, a behavior which can vary substantially within an individual’s scan, and across individuals, but which is still related to working memory [54], [55]. Using the same approach as for trait estimation, we found that median dFC model performance for estimating response time was modest (Figure 2, upper panel). Of note, the performance of shuffled data never exceeded the temporally ordered data, even at a window length of 29, which was comparable to the window length of 30 in the fIQ analysis, where shuffled FC exceeded ordered FC. Median performance of the ordered task model outperformed rest at all window lengths.

**Figure 2.**
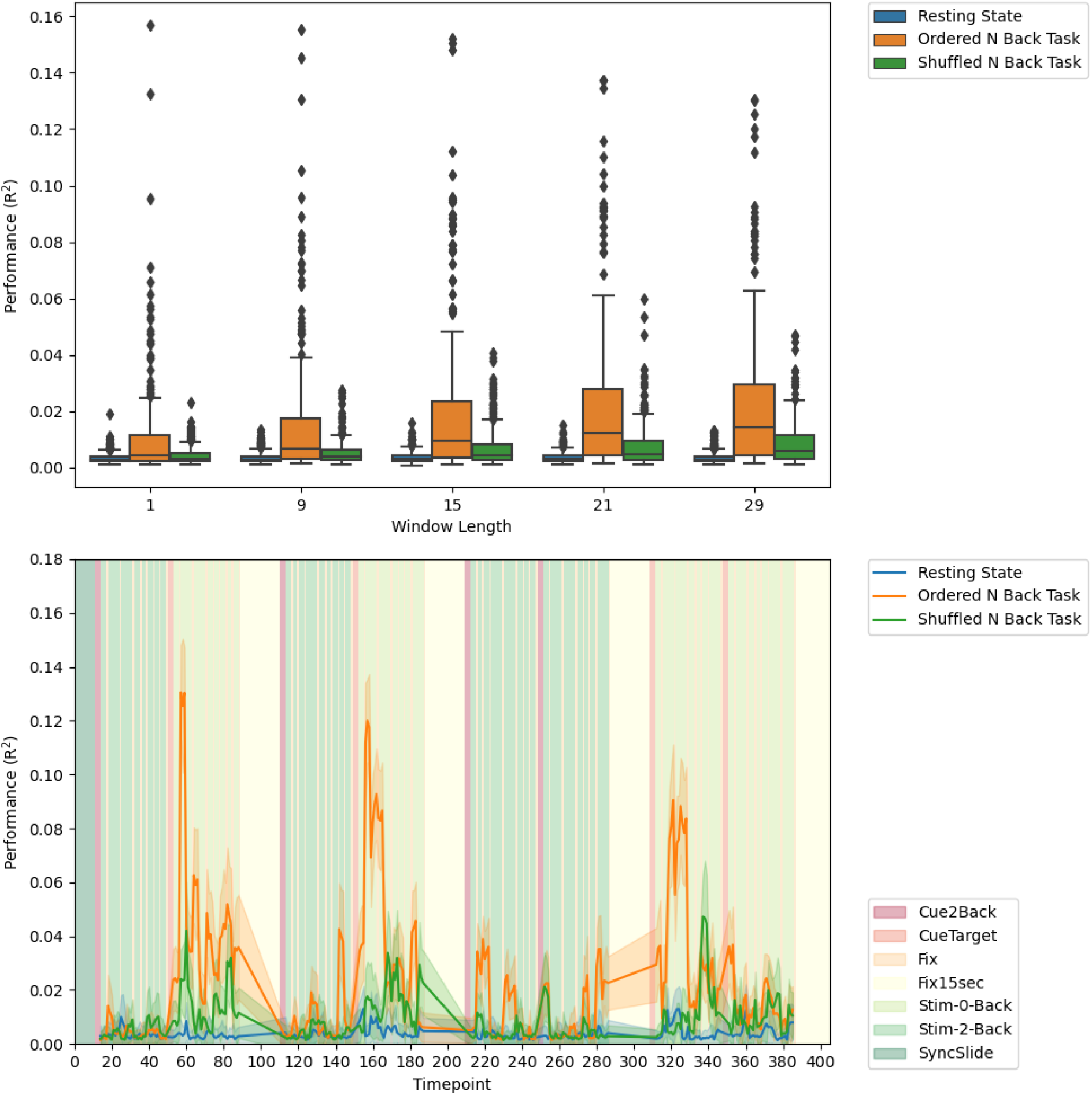
Performance of dFC models across time windows, for prediction of response time. Upper panel shows the predictive performance (y axis) of resting state dFC (blue), shuffled task dFC (green), and ordered task dFC (orange), for different window lengths (x axis). Lower panel shows the performance of each dFC type across time, for a window length of 29 timepoints only. The background reflects the task blocks, as shown in the legend in the lower right.

#### Cross validation: dFC at specific time-points

While the median performance levels across scan were relatively low, we found there was a substantial difference in prediction performance across the scan for response time. This is reflected in the number of outliers in the box plots in Figure 2, upper panel, and is shown more explicitly, for a window length of 29 timepoints (Figure 2, lower panel). At certain points in the scan, dFC could explain ~8-13% of variance in the response time of individuals. The start of the 0 back blocks appears to have been optimal, and at these points the model performance exceeded that of predicting fIQ from an equivalent amount of data, whose performance was ~3% of variance for order task data, and ~6% of variance for shuffled task data. Indeed, the dFC based prediction of response time exceeded even static FC based prediction of fIQ. In comparing different data types used in the dFC prediction, ordered task data greatly exceeded the shuffled task data, whose peak performance was around 4% variance explained. This suggests that the dFC at these specific points was maximally related to response time, compared to data from other parts of the scan. This also suggests that within scan behavior, and dFC may be highly suitable for recovering time specific spatiotemporal states. In Supplemental Figures 1-4, we replicate the plot shown in Figure 2 lower panel using each of the window lengths described in the upper panel of Figure 2 (1, 9, 15, and 21 timepoint windows). The pattern of model performance persisted across timescales, including, most notably, at the instantaneous level. As a further validation of the task ordered data prediction performance, we also used static FC to predict the response time at each trial. While we recovered a slightly similar temporal profile of model performance, the performance values were much lower, see Supplemental Figure 5.

#### In-sample replication of dFC at specific time-points

To assess the reproducibility of our findings thus far, specifically the high levels of dFC based model performance for response time, we tested how the best performing models, occurring at timepoints 59 158 and 325, performed in novel subjects in the test sample. To do so, we aggregated features and model parameters across CV folds and iterations, and generated resampled models as in previous work [50]. We applied a 90% threshold to the feature vector, that is, selecting only the features occurring in 90% or more folds across iterations. We ended up with three feature vectors, and three corresponding linear models. Again, using the CPM framework to predict response time, we applied each of the models, separately, to within sample data which had been completely omitted from the above process. We ended up with the three temporal profiles of model performance shown in Figure 3. Each of the models performed best around the timepoints they were generated, and in each case better than the other two models. This further suggests a temporal specificity to the features extracted. Altogether, these analyses suggest that when using dFC in predictive models, temporal structure matters in modelling brain-behavior relationships, and time varying behavior-based analyses are more suitable for finding temporally specific brain states than trait-based analyses.

**Figure 3.**
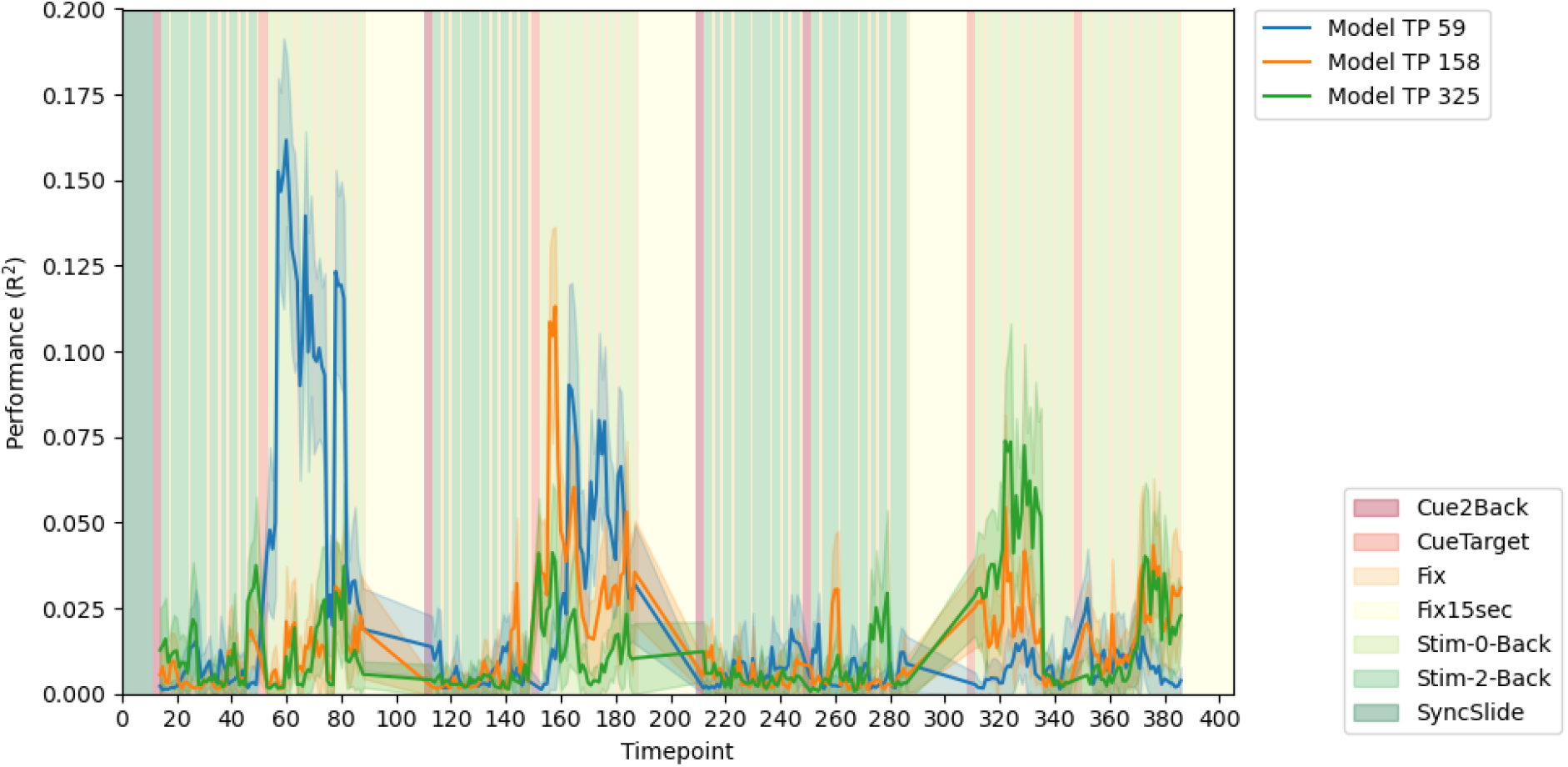
Within sample performance of resample aggregated models dFC and response time, generated at timepoints 59 (blue), 158 (orange), and 325 (green) of the N-Back HCP task. Performance is shown for features which were selected in 90% of folds across 50 iterations of split half cross validation. The background coloring reflects the task blocks, as shown in the legend in the lower right.

#### Generalizability of resampled models, within sample

#### Model Features of timepoint specific dFC models

The feature vectors used above are shown in Figure 4. Across the three models, visual, motor, limbic, and frontoparietal networks contain the greatest number of edges, consistent with task demands and previous work [56]. Model 59 has the most distributed set of edges, with many brain regions contributing to the model performance. Models 158, and 325 show increasing dependency on fewer edges, with the edges in model 325 coming mainly from four networks pairs (visual association-frontoparietal; visual association-limbic; visual network II-frontoparietal; visual network II-limbic). At a 90% threshold there are no overlapping edges between the models. A similar plot to Figure 4 is shown in Supplemental Information Figure 6, with a lower threshold (30%) to illustrate the less frequently occurring edges, where more overlap occurs. Also shown are histograms of edge occurrence frequencies in Supplemental Information Figure 7.

**Figure 4.**
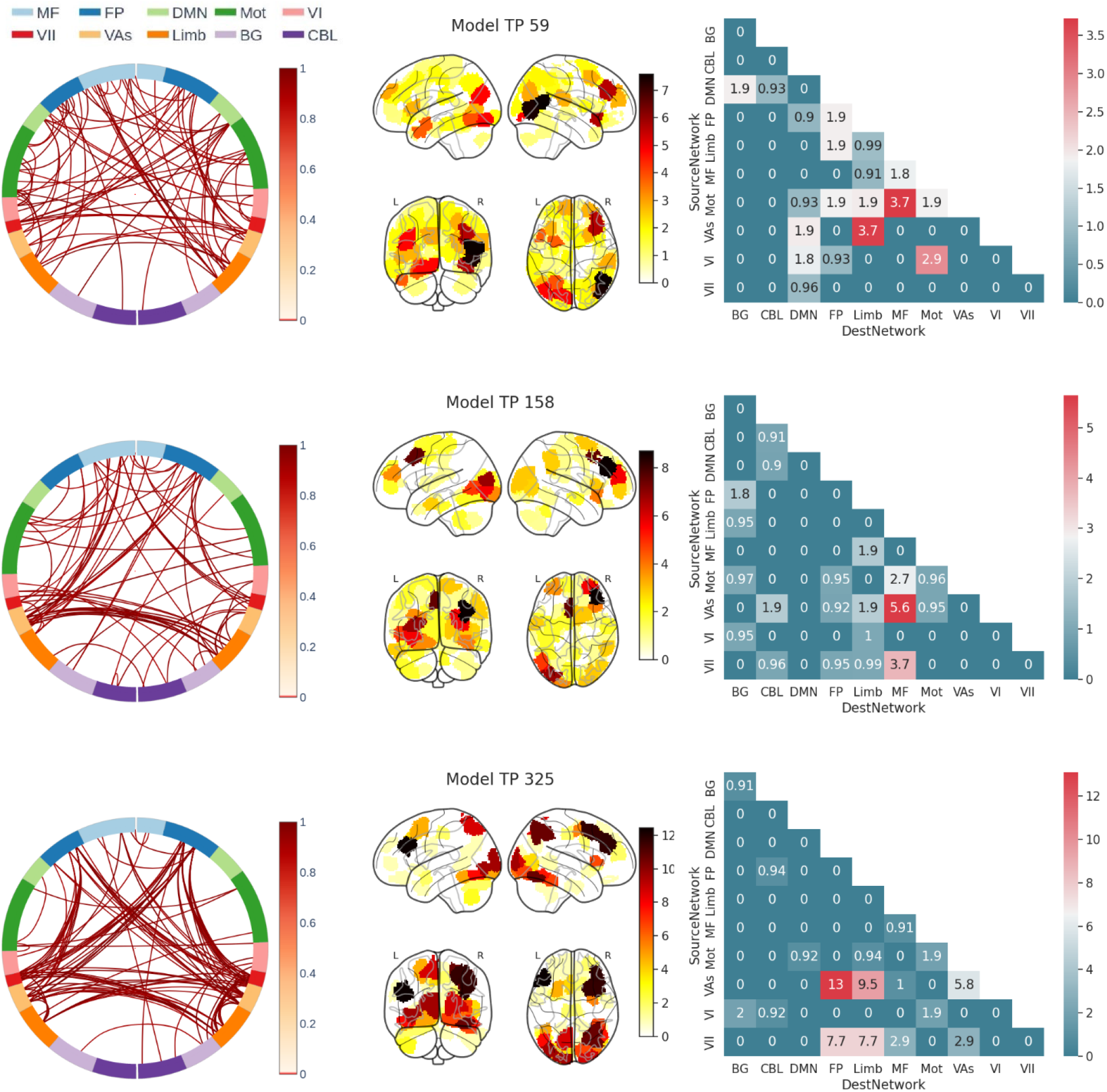
The feature sets used in Figure 3 for timepoints 59 (top), 158 (middle) and 325 (bottom). Each row shows the features for a given model, at a 90% feature frequency threshold. The first column shows a circle plot, which displays an edge-by-edge visualization of the features, weighted by occurrence. The second column shows a degree plot, which is what regions were present in the model weighted by their frequency of occurrence. The third column shows a network level matrix of what networks were represented in the feature vector, also weighted by their frequency of occurrence, and normalized for number of edges within network. The circle plot labels match the following brain networks: MF: Medio Frontal, FP: Front Parietal, DMN: Default Mode Network, Mot: Motor, VI: Visual I, VII: Visual II, VAs: Visual Association, Limb: Limbic, BG: Basal Ganglia, CBL: Cerebellum.

#### Generalizability of timepoint specific dFC models, out of sample

As a final test of model performance, the three models discussed above (from timepoints 59, 158 and 325) were applied to ABCD task data, specifically the emotional N back task (Figure 5). The ABCD scan protocols had slightly lower temporal resolution, so a window length of 25 timepoints was used to ensure similar lengths of scan time were covered. The models, which were built with HCP data only, show negligible performance. Also shown, as a point of reference, is the cross validated performance of models built from the ABCD data, using a similar method to the above. The performance of the ABCD based models does not reach those of the HCP, though they can explain 2-6% of the variance in response time at different points in the scan, again is on par with expected effect sizes of static resting state FC [53]. The pattern observed in the HCP, of high performance at the start of zero back blocks, is not replicated. As the ABCD task block ordering varied across participants, we created four groups of participants with similar task block ordering. Figure 5 shows data from group 4, groups 1-3 are shown in Supplemental Figures 8-10. Groups 1 and 3 show similar patterns to group 4, in that the models built in the HCP exhibit low performance, and with high variability across time, while the ABCD based cross validated models exhibit better performance in the range of 2-6% variance explained. Group 2 shows middling to poor performance for all models.

**Figure 5.**
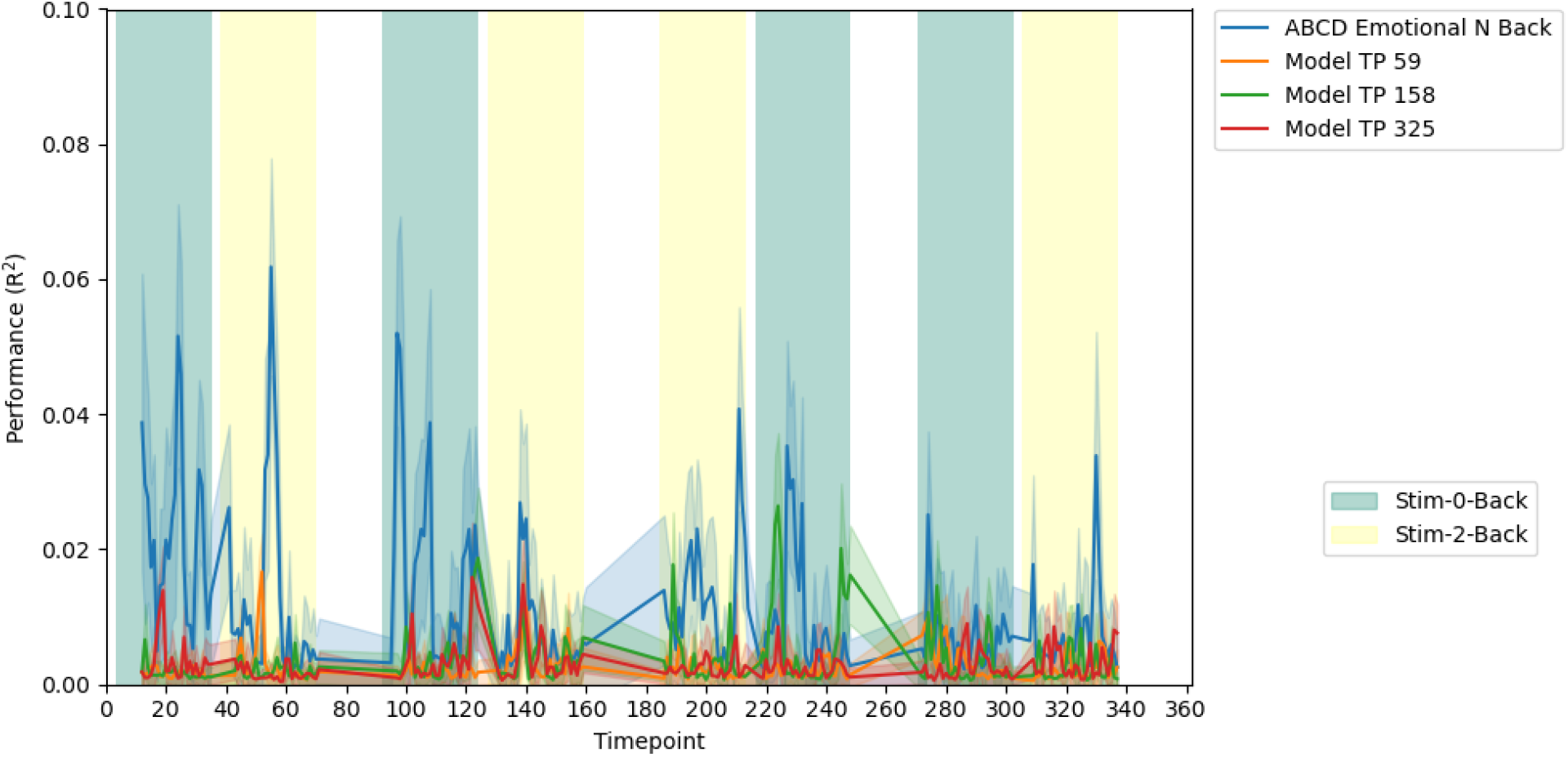
Performance of the HCP models, out of sample, (orange, green and red), juxtaposed with prediction performance of models built from ABCD data (blue). These performances are from “group 4” of the ABCD. Similar plots for the other three groups are in SI. The background reflects the task blocks, as shown in the legend in the lower right.

## Discussion

### Summary

In this study, we employ a regression model-based approach to identifying trait and behavior related brain states using phase synchrony based dFC. We contrasted rest and task-based fMRI, as well as trait and behavior, and show that task-based prediction of time-varying behavior may be more optimal for identifying temporally specific brain states. We assess the models derived using cross validation, within sample testing and out of sample testing. We found that response time to an N Back task could be optimally predicted at the start of zero back blocks, in three specific time periods, representing three different brain states with different model features, and that resampled response time models generated during these brain states replicated well within sample around the sample timepoints where they were built. This dFC-based approach for model building showed success in the ABCD sample. This framework has the potential to be useful in brain state-based investigation of time-varying behaviors in the future.

### dFC vs static FC, brain states

We examined a wide range of window lengths for fIQ prediction and a more limited range of window lengths for response time prediction. Our main dFC results for response time prediction focused on a window length of 29 timepoints, the equivalent of 20.88 seconds of data, which is at the lower end of traditional dFC analyses. However, we also saw high levels of predictive performance, for specific timepoints, at multiple temporal scales. Our initial cross validation analysis, using behavior-based prediction, was able to capture temporally specific, and predictive features at each time scale. Model performance persisted for all window lengths, including at the instantaneous level, see Supplemental Figures 1-4. It was also possible to predict fIQ from short time dFC, though the performance levels never exceeded those of static FC, and the predictive features tended to be more similar across the scan.

dFC allows for more temporally precise measurement of FC, with the possibility of investigating temporally specific aspects of functional brain connectivity during scanning. Traditionally dFC based studies of brain states have used longer time windows to estimate connectivity, such as 40-100 seconds [57], [58], often with longer repetition time than the protocols used in this study. There are studies which have shorter window sizes [10], [13], or no window at all [27]. However, these studies still aggregate data from the whole scan into the derivation of brain states. Our regression-based approach, based on phase synchrony dFC, can go beyond more typical sliding window methods, and allow for estimation of brain states using very little data per participant, down to the level of individual timepoints. When combined with short TR acquisitions, this allows for high temporal specificity.

There is precedent for predicting behavioral and perceptual outcomes from short time FC, though without the explicit focus on brain states. An early study in the field by Thompson et al. [14] modelled response time to a psychomotor vigilance task using regression,[14] though it was only based on short time connectivity estimates from two brain networks, and used a relatively small sample size. Classification approaches have also been used to predict auditory [59] and tactile stimuli [60], as well as preference for faces [61]. Later, Kucyi et al. showed that “response times” to a self-paced task, characterized by repetitive button pressing, could be used to determine periods of fatigue or distraction [62]. Working memory specific reconfiguration of dFC has also been well documented [63]–[65]. This study is well placed among these findings, given that an individual’s response time to an N back task likely incorporates many phenomena, including perception, attention and fatigue. Our study was unique in that it combined a regression approach, an order of magnitude more subjects than some of the studies listed above, a parcellation scheme with whole brain coverage, and only ~20 seconds of data or less per participant for any given model. We assert that the method detailed here is relevant both to the study of behaviorally related brain states and for short time predictive modelling.

### Resting state vs task, Nulls

As discussed, we sought to find temporally specific spatial patterns of brain connectivity which were related to a trait or behavior of interest. In order to assess the utility of task based dFC, we compared the predictive power of (a) temporally ordered task-based fMRI, (b) temporally shuffled task-based fMRI, and (c) resting state fMRI. Previously, it has been shown that static resting state FC is a poorer predictor of fIQ than task-based connectivity [44], though it was unclear if this would hold for dynamic connectivity, or for response time prediction. Here, we confirm that resting state connectivity is indeed a poorer predictor of fIQ at the dynamic level, and its predictive value remains negligible until 4-5 mins of data are included. A similar trend is seen for response time. We suggest that resting state fMRI, within the context of dynamic assessment of behaviorally related brain states, can provide a useful foil for task based dFC, serving as a null space for comparisons. Indeed, it has the same nuisance factors as task-based data, while the cognitive content is more spontaneous and uncoordinated [66], [67]. Another comparison conducted was whether temporally ordered task data led to better predictive models than randomly ordered, or shuffled, task data. We found that shuffled task data performed better for trait prediction, while temporally ordered data worked better for behavior prediction. This suggests that for trait-based predictions, drawing data from across the scan is advantageous, whereas, for behavior, more precise estimates of connectivity are better. Cross subject synchrony, or lack thereof, is an important aspect in FC based analyses, and synchrony and desynchrony may reflect different, but equally cognitively relevant processes [68].

As well as wanting to assess the biological relevance of time ordered task data, we also sought to bolster our confidence in the results of our analytic methods. As discussed in the introduction, a common method in dFC research is the construction of null spaces, simulating random data from real covariance matrices and power spectra [29], [30]. The goal of these efforts is to interrogate the validity of the results drawn from real fMRI data, in particular resting state data. However, it is difficult to fully replicate real physiological noise, and motion, present in these data. We used resting state dFC as our “null”. Task-based dFC analyses, which rely on cross subject synchrony, can be juxtaposed with a resting dFC to provide a rigorous statistical comparison. This approach may prove advantageous in harnessing dFC for use in a clinical setting where a particular trait or behavior of interest is be interrogated with a task paradigm can be interrogated with a task paradigm while using the same subject’s resting-state data as the null condition.

### Modelling and Generalizability

We derived three regression models of time varying behavior, which replicated within sample, on unseen subjects. The resampling technique used to generate mean feature vectors allowed us to assess how much variance there is in feature selection within sample. Boiling down feature sets to the most salient features, that is the ones which occur most frequently, and assessing on unseen data offers a path forward to interrogating and replicating the success of brain behavior models [50]. Generalizability is a key roadblock in the development of brain behavior models, with overfitting the norm rather than the exception. The evaluation of models in out of sample data can be difficult. Failure to replicate within sample would indicate complete overfitting, but one can interpret a lack of generalization to other datasets in different ways. This topic has been covered previously, including in our own work [50], [69]. It is possible that a model can capture salient features reflecting underlying cognition, while failing to generalize. The ABCD data comes from a multisite on-going initiative, presenting possible confounds that may arise due to multiple scanner and site differences [70]. This is in addition to sample based phenotypic differences, age being a prime factor which impacts working memory performance [71], and task paradigm differences. The N Back was used in both HCP and ABCD, but the ABCD used different stimuli, designed to evoke emotional responses, in addition to testing the working memory of participants [38]. Each of these factors could impede generalization across these datasets. It may be that case that the cognition underlying response time to a working memory N back task in adults, as represented by dFC, is fundamentally different to response time to an emotional N back task in 9–11-year-olds. That is to say, perhaps the dataset we chose to attempt generalization in was not the ideal test case. Nonetheless, it is imperative to assess the degree to which model generalization is impeded and attempt to overcome these differences. In that vein, we hypothesized that by reducing the amount of data taken from a given dataset, we may be able to capture only the most relevant parts of a study, potentially aiding generalizability. Thus, while between study differences may have contributed to a lack of transfer of models between studies, the ABCD data did validate our approach to model building, as we could generate within sample models in both data sets, which explained variance in response in each independent dataset.

### Trait vs Behavior

So far, we have focused on the imaging aspect of the brain behavior modelling. The terms trait and behavior are often used interchangeably; in this case we consider traits to be either unchanging, or changing over much longer periods of time, when compared to time-varying behavior which can change moment-to-moment [72]. Often traits are used as axes of contrast across participants in dFC analyses, for example a diagnosis, or a score on a cognitive assessment. Aside from the fact that these are not permanent attributes, and may vary across different periods of time, they do not vary over the length of scan. We would argue that this makes them poor candidates for brain state-based dFC analyses. A better approach may be to incorporate a manipulatable behavior, related to the trait under consideration. As mentioned above, reproducibility and generalization should be of major concern in any study. A recent paper by Elliot et al proposes several methods for conducting better studies including i) aggregating more repeated in person measurements, ii) modelling points of the scan with stable variability, iii) better accounting of physiological noise and iv) designing more reliable tasks [73]. The framework described above can be used to address points ii) and iv) by identifying stable brain states which have sustained relationships with behaviors of interested, as assessed via continued high performance of the model. This can then be used to probe the usefulness of a task fMRI paradigm at the maximum allowable temporal resolution. This opens up the possibility of a feedback loop between task design and model performance, whereby tasks can be designed to maximize model performance, or effect size. Maximizing the effect size for the studied relationship is particularly relevant in the context how effect size and sample size affect the power, and therefore reliability, of a study [53], [74]–[76].

## Conclusion

We have developed a framework for identifying behaviorally relevant short time brain states, as represented by dynamic functional connectivity maps. We have developed a framework for identifying behaviorally relevant short time brain states, as represented by dynamic functional connectivity maps. We find that drawing data from across the scan is better for trait-based prediction, but more isolated and temporally specific parts of the scan are better for developing models that predict in-task performance. The resample aggregated dynamic functional connectivity models of behavior replicated within sample using unseen HCP data, and this same modeling framework showed success in the estimating variance behavior in the ABCD dataset. Future work could assess if the framework described here could be used to model other time-varying behaviors under different task conditions.

## Acknowledgements

CH was supported by a Medical Scientist Training Program training grant (NIH/NIGMS T32GM007205).

Data were provided in part by the Human Connectome Project, WU-Minn Consortium (principal investigators, D. Van Essen and K. Ugurbil; 1U54MH091657) funded by the 16 US National Institutes of Health (NIH) institutes and centers that support the NIH Blueprint for Neuroscience Research; and by the McDonnell Center for Systems Neuroscience at Washington University.

Data used in the preparation of this article were obtained from the Adolescent Brain Cognitive Development (ABCD) Study [38], held in the NIMH Data Archive (NDA). This is a multisite, longitudinal study designed to recruit more than 10,000 children age 9-10 and follow them over 10 years into early adulthood. The ABCD Study^®^ is supported by the National Institutes of Health and additional federal partners under award numbers U01DA041022, U01DA041028, U01DA041048, U01DA041089, U01DA041106, U01DA041117, U01DA041120, U01DA041134, U01DA041148, U01DA041156, U01DA041174, U24DA041123, and U24DA041147. A full list of supporters is available at abcdstudy.org/nih-collaborators. A listing of participating sites and a complete listing of the study investigators can be found at abcdstudy.org/principal-investigators.html. ABCD consortium investigators designed and implemented the study and/or provided data but did not necessarily participate in analysis or writing of this report. This manuscript reflects the views of the authors and may not reflect the opinions or views of the NIH or ABCD consortium investigators. The ABCD data repository grows and changes over time. The ABCD data used in this report came from NIMH Data Archive Digital Object Identifier 10.15154/1504041. DOIs can be found at nda.nih.gov/study.html?id=721.

## Data and Code Availability

The HCP data that support the findings of this study are publicly available on the ConnectomeDB database (https://db.humanconnectome.org). The ABCD data used in this report came from NIMH Data Archive Digital Object Identifier 10.15154/1504041. DOIs can be found at nda.nih.gov/study.html?id=721. Code for conducting the analyses described here can be found at https://github.com/YaleMRRC/dFCAnalysis

