## Supplemental Information for "Identifying dynamic reproducible brain states using a predictive modelling approach"

### Temporal performance of response time dFC models

#### *Instantaneous dFC prediction of response time*

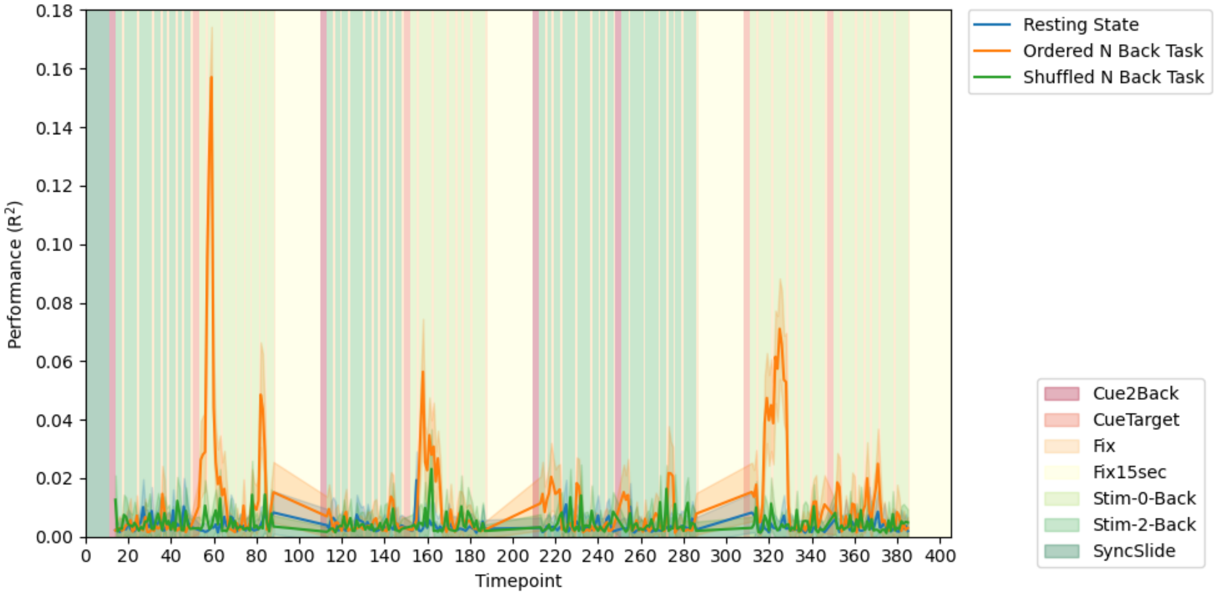


Figure 1 - The predictive performance (y axis) of different models based on resting state dFC (blue), shuffled task dFC (green), and ordered task dFC (orange), for prediction of N back response time. The window length for dFC calculation was 1 timepoint. The background reflects the task blocks, as shown in the legend in the lower right.

#### *9 timepoint windowed dFC prediction of response time*

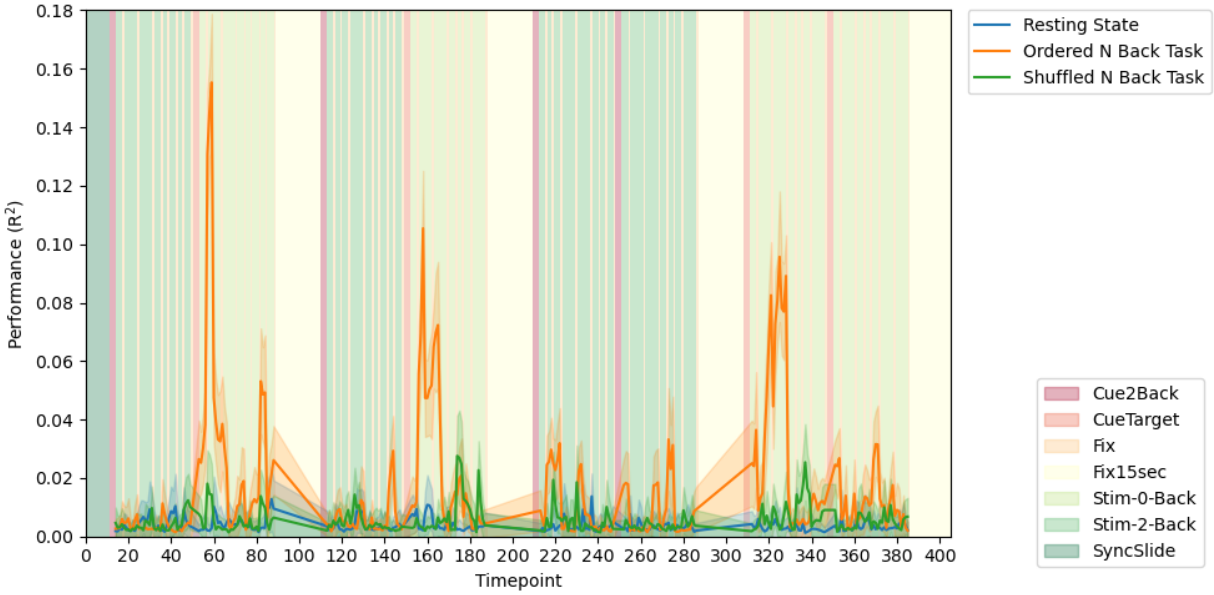


Figure 2 - The predictive performance (y axis) of different models based on resting state dFC (blue), shuffled task dFC (green), and ordered task dFC (orange), for prediction of N back response time. The window length for dFC calculation was 9 timepoints. The background reflects the task blocks, as shown in the legend in the lower right.

### *15 timepoint windowed dFC prediction of response time*

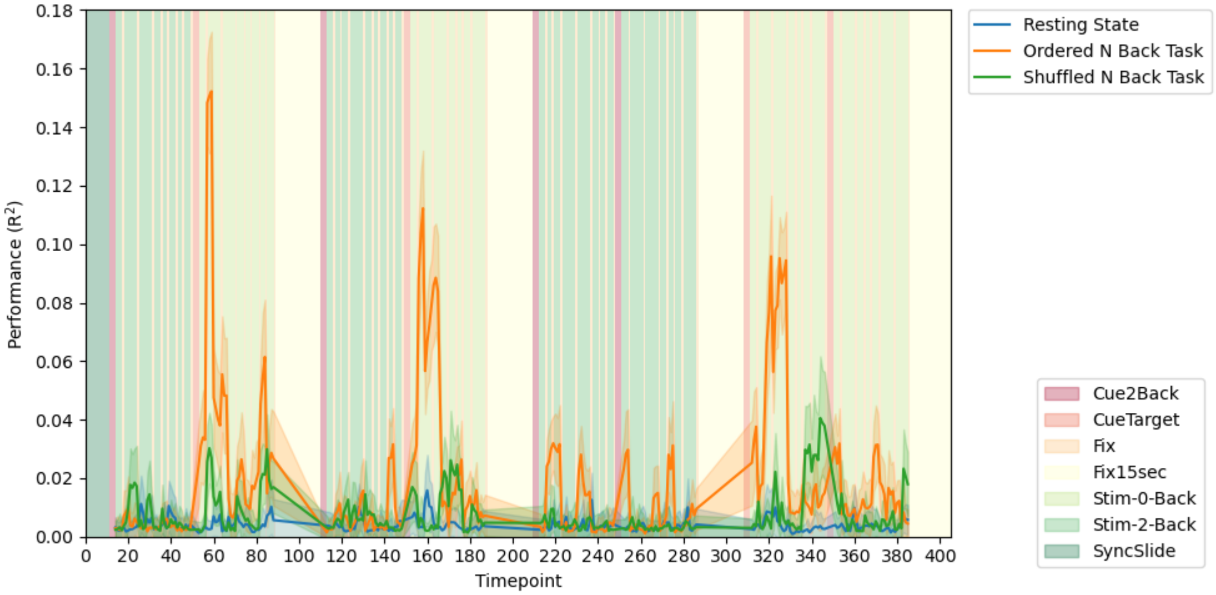


Figure 3 - The predictive performance (y axis) of different models based on resting state dFC (blue), shuffled task dFC (green), and ordered task dFC (orange), for prediction of N back response time. The window length for dFC calculation was 15 timepoints. The background reflects the task blocks, as shown in the legend in the lower right.

### *21 timepoint windowed dFC prediction of response time*

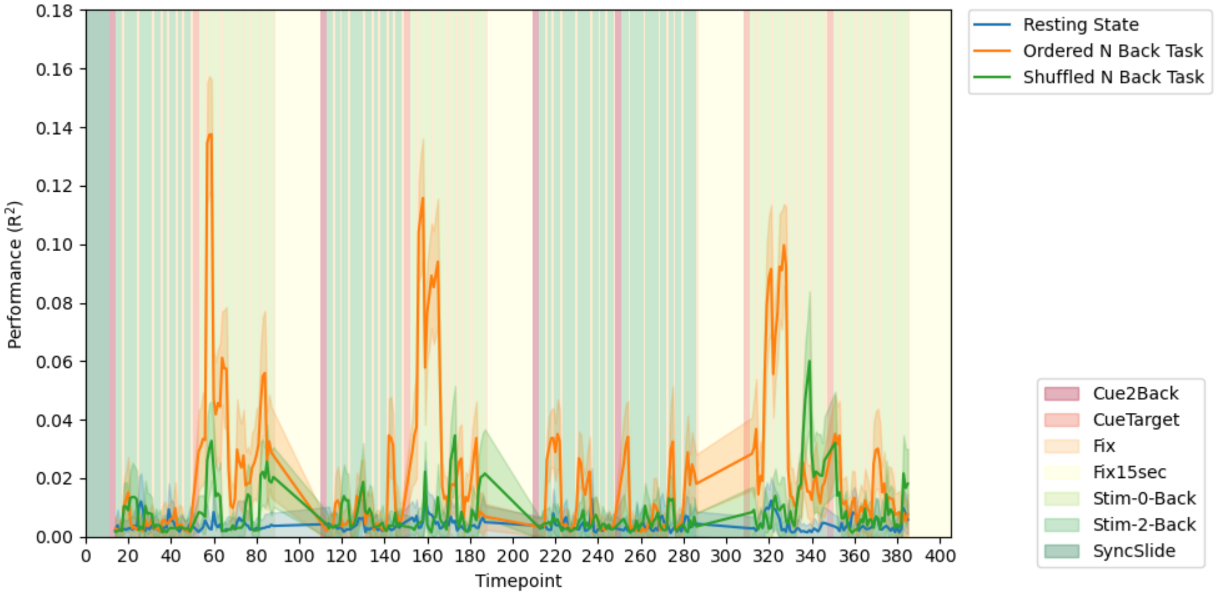


Figure 4 - The predictive performance (y axis) of different models based on resting state dFC (blue), shuffled task dFC (green), and ordered task dFC (orange), for prediction of N back response time. The window length for dFC calculation was 21 timepoints. The background reflects the task blocks, as shown in the legend in the lower right.

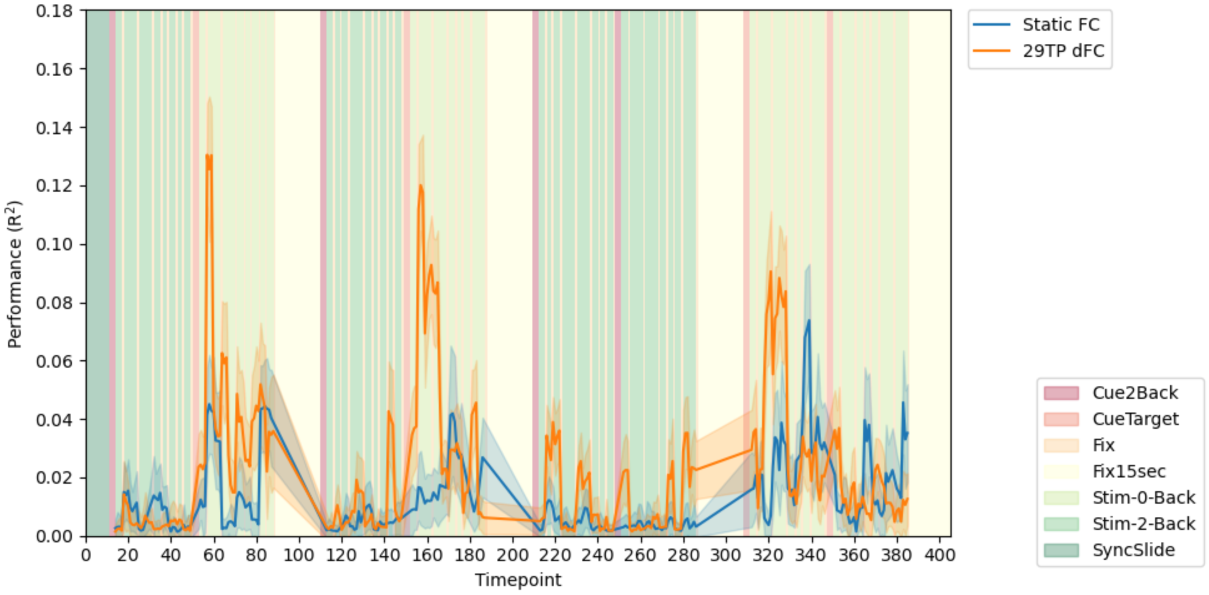


Figure 5 - The predictive performance (y axis) of different models based on static FC (blue), and ordered task dFC (orange), for prediction of N back response time. The window length for dFC calculation was 29 timepoints, and the full scan length was used for calculating static FC. The background reflects the task blocks, as shown in the legend in the lower right.

### Model features selected

#### *Edge plots with 30% feature threshold*

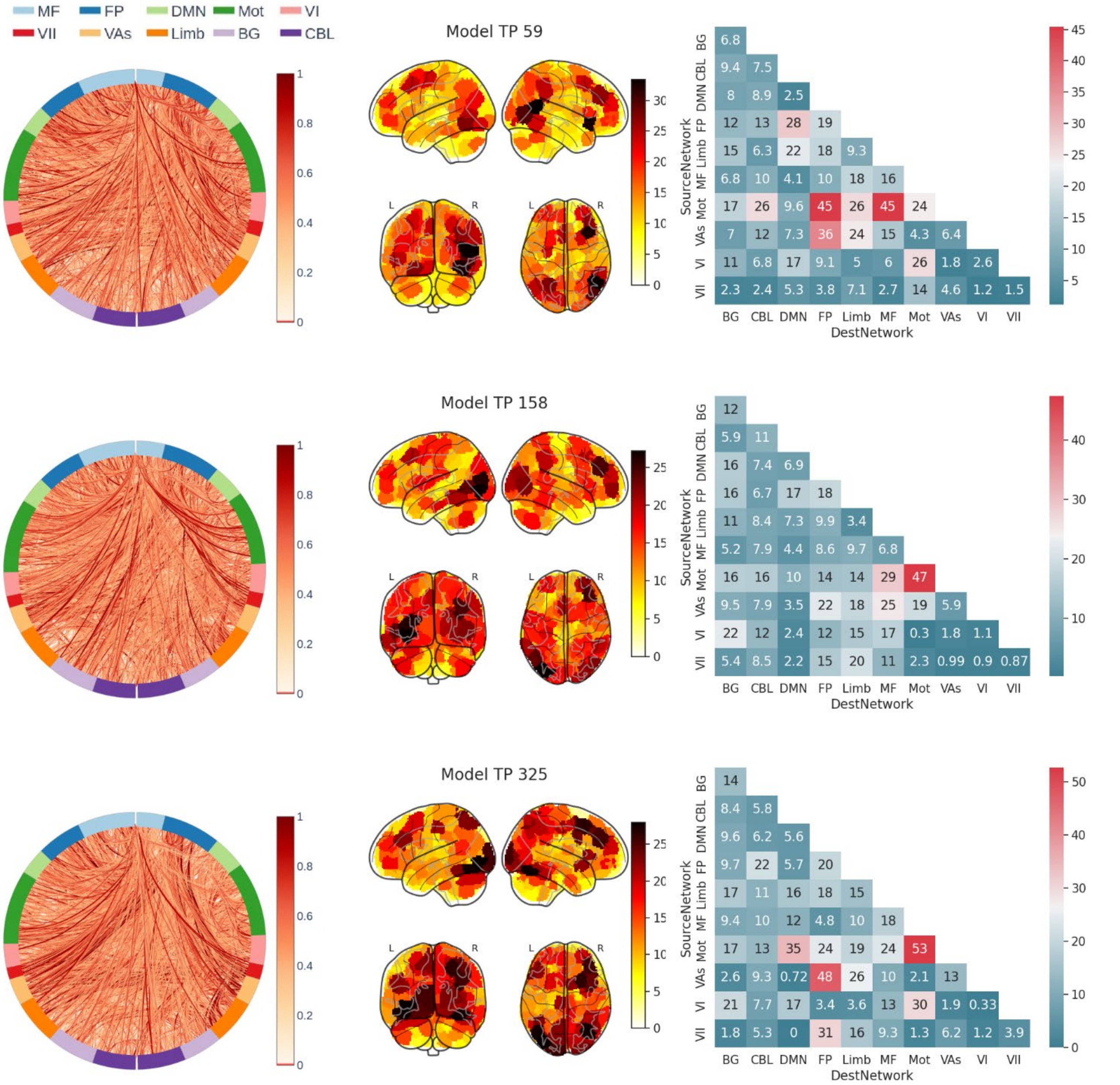


Figure 6 – The feature sets used in Figure 3 for timepoints 59 (top), 158 (middle) and 325 (bottom). Each row shows the features for a given model, at a 30% feature frequency threshold. The first column shows a circle plot, which displays an edge-by-edge visualization of the features, weighted by occurrence. The second column shows a degree plot, which is what regions were present in the model weighted by their frequency of occurrence. The third column shows a network level matrix of what networks were represented in the feature vector, also weighted by their frequency of occurrence, and normalized for number of edges within network. The circle plot labels match the following brain networks: MF: Medio Frontal, FP: Front Parietal, DMN: Default Mode Network, Mot: Motor, VI: Visual I, VII: Visual II, VAs: Visual Association, Limb: Limbic, BG: Basal Ganglia, CBL: Cerebellum.

*Distribution of edges selected for three models used*

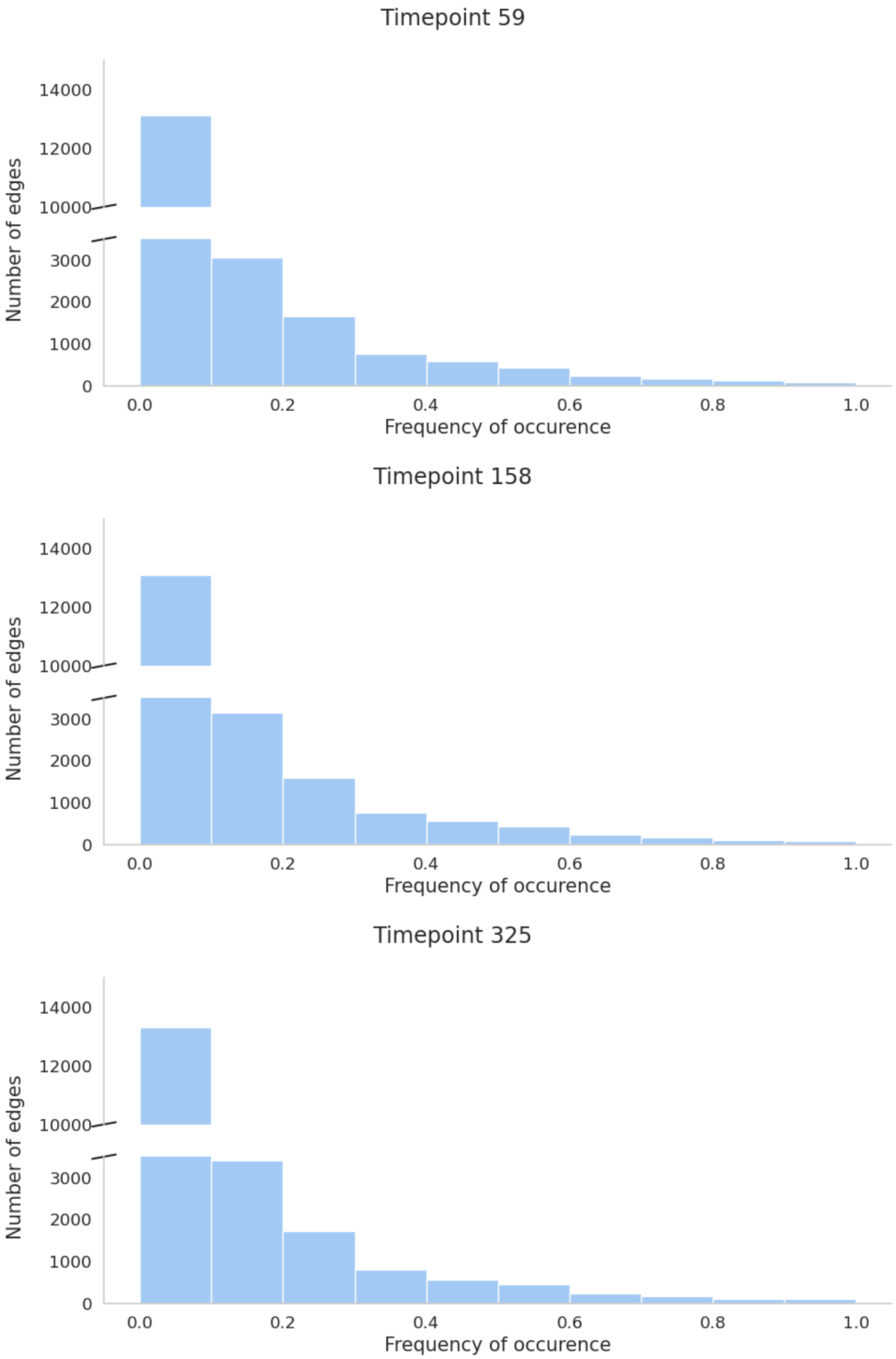


Figure 7 – The distribution of edge frequency occurences for each selected dFC model of N back response time. The top panel is for the model generated from timepoint 59, the second row for timepoint 158 and the third for timepont 325.

### Temporal performance of ABCD Cross Validation and out of sample HCP Performance

#### *Group 1*

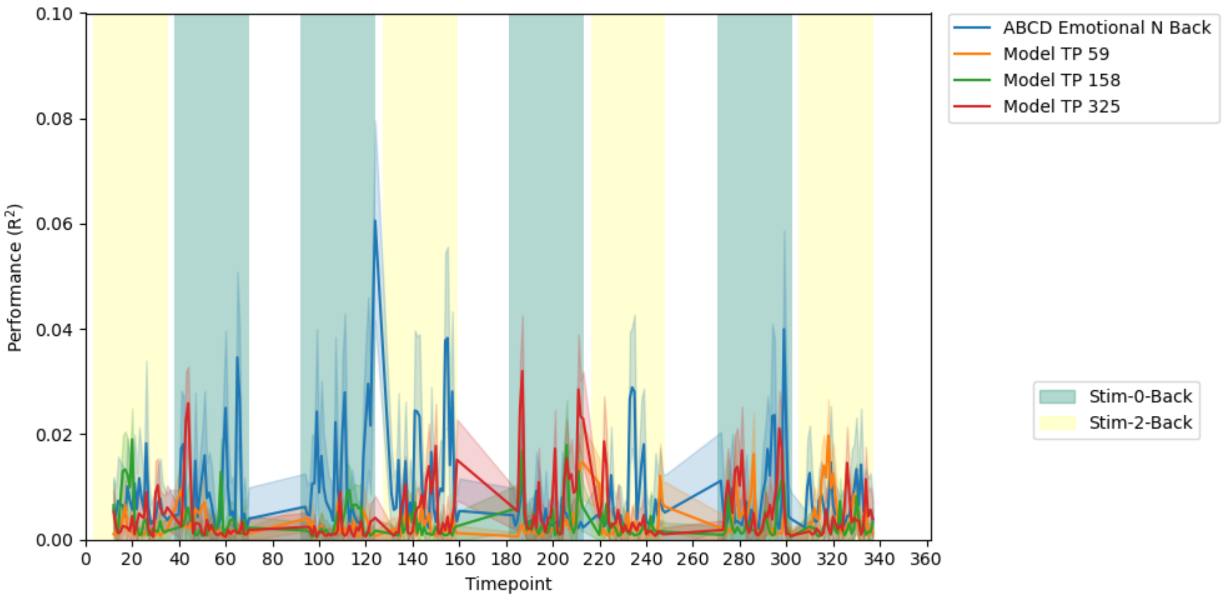


Figure 8 – Performance of the HCP models, out of sample, (orange, green and red), juxtaposed with prediction performance of models built from ABCD data (blue). These performances are from “group 1” of the ABCD. Similar plots for the other three groups are in SI. The background reflects the task blocks, as shown in the legend in the lower right.

#### *Group 2*

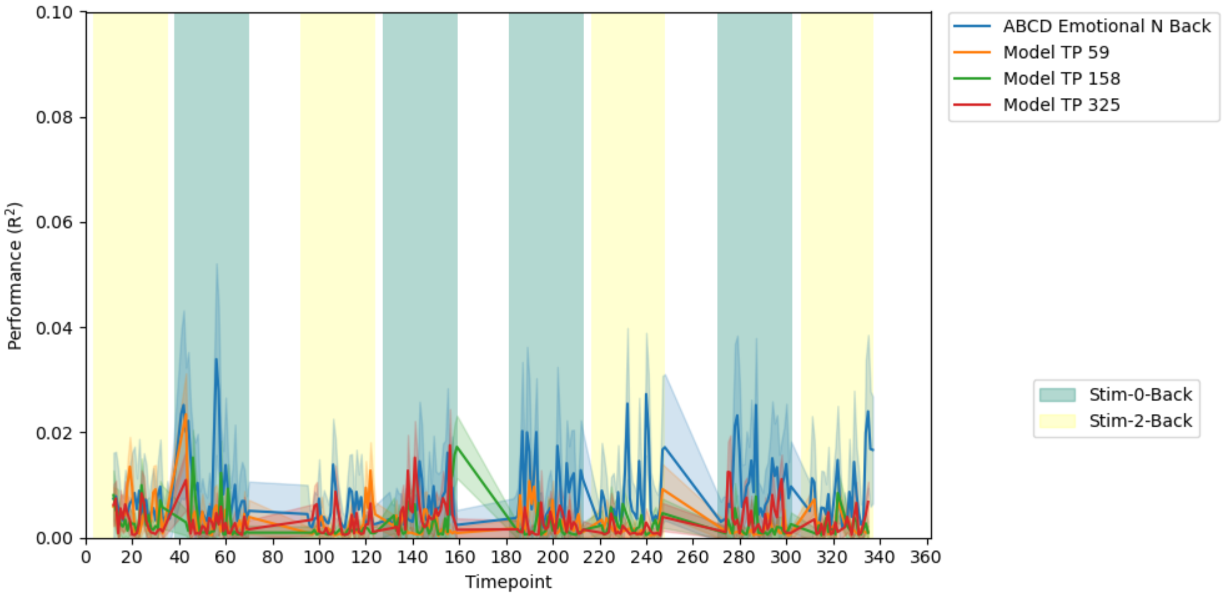


Figure 9 – Performance of the HCP models, out of sample, (orange, green and red), juxtaposed with prediction performance of models built from ABCD data (blue). These performances are from “group 2” of the ABCD. Similar plots for the other three groups are in SI. The background reflects the task blocks, as shown in the legend in the lower right.

#### *Group 3*

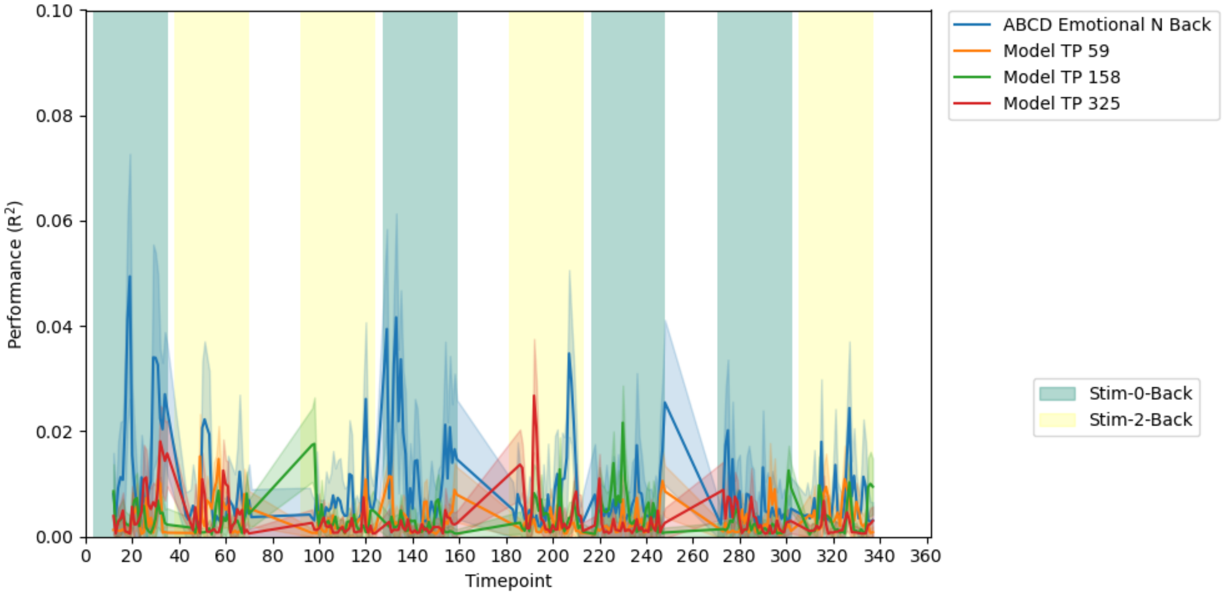


Figure 10– Performance of the HCP models, out of sample, (orange, green and red), juxtaposed with prediction performance of models built from ABCD data (blue). These performances are from “group 3” of the ABCD. Similar plots for the other three groups are in SI. The background reflects the task blocks, as shown in the legend in the lower right.
